# SARS-CoV-2 infection of human brain microvascular endothelial cells leads to inflammatory activation through NF-κB non-canonical pathway and mitochondrial remodeling

**DOI:** 10.1101/2022.06.16.496324

**Authors:** Silvia Torices, Carolline Soares Motta, Barbara Gomes da Rosa, Anne Caroline Marcos, Liandra Alvarez-Rosa, Michele Siqueira, Thaidy Moreno-Rodriguez, Aline Matos, Braulia Caetano, Jessica Martins, Luis Gladulich, Erick Loiola, Olivia RM Bagshaw, Jeffrey A. Stuart, Marilda M. Siqueira, Joice Stipursky, Michal Toborek, Daniel Adesse

**Author notes:** **Corresponding authors: Daniel Adesse**, Laboratório de Biologia Estrutural, Instituto Oswaldo Cruz, Fiocruz, Avenida Brasil, 4365, Pavilhão Carlos Chagas, sl 307, Rio de Janeiro, RJ 21040-360, Brazil, **Joice Stipurksy**, Laboratório Compartilhado, Instituto de Ciências Biomédicas, Universidade Federal do Rio de Janeiro, Avenida Carlos Chagas Filho. Equally contributed as senior authors (Michal Toborek, Daniel Adesse). Equally contributed as first authors (Silvia Torices, Carolline Soares Motta).

## Abstract

Neurological effects of COVID-19 and long-COVID-19 as well as neuroinvasion by SARS-CoV-2 still pose several questions and are of both clinical and scientific relevance. We described the cellular and molecular effects of the human brain microvascular endothelial cells (HBMECs) *in vitro* infection by SARS-CoV-2 to understand the underlying mechanisms of viral transmigration through the Blood-Brain Barrier. Despite the low to non-productive viral replication, SARS-CoV-2-infected cultures displayed increased apoptotic cell death and tight junction protein expression and immunolocalization. Transcriptomic profiling of infected cultures revealed endothelial activation via NF-κB non-canonical pathway, including RELB overexpression, and mitochondrial dysfunction. Additionally, SARS-CoV-2 led to altered secretion of key angiogenic factors and to significant changes in mitochondrial dynamics, with increased mitofusin-2 expression and increased mitochondrial networks. Endothelial activation and remodeling can further contribute to neuroinflammatory processes and lead to further BBB permeability in COVID-19.

## INTRODUCTION

Coronavirus disease 2019 (COVID-19), caused by infection with severe acute respiratory syndrome-related coronavirus 2 (SARS-CoV-2) remains at major health threat globally. The U.S. continues to lead the world with a total of 47.9 million COVID-19 cases and 773,887 deaths at the end of November 2021 (WHO). Even though vaccines, which mostly prevent serious illness and death, have been widely available in the U.S. and many countries, only 59% of the overall population are fully vaccinated. This is below the estimated 85-90% threshold assumed to be needed to stop the spread of SARS-CoV-2 and make the virus endemic. Globally, COVID-19 cases continue rising, and new variants and subvariants (e.g. omicron BA.1, BA.2 [BA.2.12, BA.2.12.1]) were recently identified in different countries, spreading globally (WHO, 2022).

SARS-CoV-2 is a member of the family of β-coronaviruses, similar to two other highly pathogenic coronaviruses: severe acute respiratory syndrome coronavirus (SARS-CoV) and Middle East respiratory syndrome coronavirus (MERS-CoV). Initial SARS-CoV-2 infection cases investigated led to the isolation of the virus in human respiratory epithelial cells (CDC; Zhu et al., 2020) and its genome sequencing deposited (GISAID accession IDs: EPI_ISL_402119, 402120 and 402121). SARS-CoV-2 is an enveloped, positive-sense, and single-stranded RNA virus. Its genome encodes non-structural proteins (such as 3-chymotrypsin-like protease, papain-like protease, helicase, and RNA-dependent RNA polymerase; all key enzymes in the viral life cycle), structural proteins (spike [S] protein, membrane [M] protein, envelope [E] protein and nucleocapsid [N] protein) and accessory proteins.

It is now known that SARS-CoV-2 infects human cells through the ligation of the S1 subunit of the S protein with host cell receptors, especially the angiotensin-2 converting enzyme (ACE2), that serves as an entry receptor to the virus, representing its main route of entry into the host cell (Wang et al., 2020). Pulmonary, cardiac and intestinal epithelia and endothelial cells express high levels of ACE-2 (CDC, 2020). Upon S1-ACE2 interaction, a transmembrane serine protease 2 (TMPRSS2) is required for priming of the S protein and viral entry into the cell (Coronaviridae Study Group of International Committee on Taxonomy of Viruses, 2020; Zhu et al., 2020; Ni et al., 2019). Along with ACE2 and TMPRSS2, several other proteins have been suggested to participate in SARS-CoV-2 entry into human cells, such as ADAM metallopeptidase domain 17 (ADAM17) (Torices et al, 2021; Hoffmann et al., 2020), dipeptidyl peptidase 4 (DPP4) (Zhou et al., 2020; Shulla et al., 2011), angiotensin II receptor type 2 (AGTR2) (Zipeto et al., 2020; Schreiber et al., 2020), basigin (BSG, also called extracellular matrix metalloproteinase inducer [EMMPRIN] or cluster of differentiation 147 [CD147]) (Solerte et al., 2020; Bassendine et al, 2020), aminopeptidase N (ANPEP) (Cui et al., 2020) and cathepsin B/L (de Souza et al. 2020; Wang et al., 2020).

Among the symptoms most commonly observed in COVID-19 patients, alterations of neural functions are frequently detected, from mild cases with loss of taste and smell, dizziness and headaches, to more extreme cases with occurrence of acute cerebrovascular disease, including episodes of vascular encephalic accidents, loss of consciousness, ataxia and epilepsy (Qiao et al., 2020). The Central Nervous System (CNS)-related symptoms are also prominent for so called chronic or long-COVID. The CNS is a well-documented target of β-coronavirus infections, such as SARS-CoV-2, and to date several studies detected SARS-CoV-2 in the brain and the cerebrospinal fluid of COVID-19 patients (Qi et al., 2020; Padmanabhan et al., 2020; Istlfl et al., 2020; Ellul et al., 2020). Distinct routes of SARS-CoV-2 entry into the brain have been proposed, such as the olfactory nerve (Mao et al., 2020; Solomon et al., 2020; Virhammar et al., 2020), the choroid plexus and the blood-brain barrier (BBB) (Baig et al., 2020). However, the contribution of the BBB may be particularly important due to presence of the virus in the bloodstream allowing the passage of viral particles through the wall of brain capillaries to brain parenchyma. While the mechanisms of SARS-CoV-2 neuroinvasion are not fully understood, it has been suggested that infection of BBB capillaries-composing cells is critical to trigger CNS impairment (Li et al., 2020a; Li et al., 2020b). Several essential characteristics that ensure BBB function are supplied by endothelial cells that express specific types of tight junction proteins (Briguglio et al., 2020; Alam et al., 2020; Achar et al., 2020).

Several reports have correlated the infection outcome with vascular dysfunction, establishing vascular inflammation and cytokine storms promoted by immune responses as critical factors contributing to worsening of the clinical condition and even death. Endothelial dysfunction may have important consequences, which include ischemia, altered angiogenesis and coagulation, inflammation, and tissue edema. Therefore, COVID-19-related endothelitis could explain the systemic microcirculatory dysfunction observed in patients, including chronic form of this disease. It was recently demonstrated that the treatment of human brain microvascular endothelial cells with recombinant S1 protein resulted in endothelial permeability and altered the levels of pro-inflammatory cytokines (Buzhdygan et al., 2020). However, little is known about the involvement of brain microvasculature in brain infection by SARS-CoV-2, which may result in endothelial activation and hyper-inflammatory responses. Even less is known if damages to the BBB could be propagated to neural tissue, and therefore be the triggering mechanism of neural abnormalities that promote neurological symptoms observed in COVID-19 patients.

In the present work we describe cellular and molecular effects of HBMEC infection by SARS-CoV-2 in order to gain insight into possible routes by which the virus affects the BBB and invades the brain parenchyma. Transcriptomic analyses revealed activation of noncanonical NF-κB signaling pathway and changes in mitochondrial quality control, which combined could induce endothelial activation and promote increased neuroinflammation in Neuro-COVID-19.

## METHODS

### 1. Cell culture

Human brain endothelial cells (HBMECs) were a gift from Prof. Dennis Grab (Department of Pathology, Johns Hopkins School of Medicine). Cells were immortalized using a SV40-LT plasmid (Stins et al., 2001) and were maintained in 199 medium with 10% fetal bovine serum (FBS) and 1% antibiotics (penicillin/streptomycin, ThermoFisher) up to passage 38. Vero E6 cells (African green monkey kidney epithelial cells) were used as gold standard for viral isolation and propagation and were used in a few experiments as a positive control for efficient SARS-CoV-2 infection. Vero E6 cells culture medium consisted of Dulbecco’s Modified Eagle Medium (DMEM) formulated with D-glucose (4.5 g/l) and L-Glutamine (3.9 mM) supplemented with 100× penicillin-streptomycin solution (to final 100 U/ml and 100 μg/ml, respectively) and with inactivated FBS (USDA-qualified region FBS) at 10%. Both cell and viral cultures were incubated at 37°C and 5% CO_2_.

### 2. SARS-CoV-2 isolate

All the procedures associated to the viral culture and further infection assays were performed in biosafety level 3 laboratory, in accordance with the WHO guidelines (WHO, 2021). The SARS-CoV-2 isolate used in this study was previously obtained from a respiratory sample collected from a COVID-19 patient diagnosed in March 2020, in Brazil, as part of the Brazilian Ministry of Health surveillance system. Viral isolation and genetic characterization were previously described (Matos et al, 2022). All procedures involving patient samples were approved by the Committee of Ethics in Human Research of the Oswaldo Cruz Institute (registration number CAAE 68118417.6.0000.5248).

### 3. SARS-CoV-2 infection

Cells were cultured to obtain confluent monolayers. After that, cells were washed once with PBS and SARS-CoV-2 inoculums corresponding to indicated multiplicities of infection (MOI) were incubated for one hour. After infection, inoculums were removed from cells and replaced by their appropriate supplemented medium with TPCK treated trypsin at 1 μg/mL.

### 4. Viral quantification

We evaluated SARS-CoV-2 replication of infected cells over time by measuring the number of viral RNA copies in their culture media over time. Viral RNA was extracted from 140 μL of cell-free culture media using QIAamp Viral RNA mini kit (Qiagen, Germany), according to manufacturer’s instructions. Reverse transcription and SARS-CoV-2 gene amplification were performed in one-step reactions with a quantitative real time PCR kit developed by Biomanguinhos Institute (Fiocruz, Brazil), in an ABI 7500 thermocycler (Applied Biosystems, USA). As a quantification standard, we used a SARS-CoV-2 plasmid control containing the reference sequence of viral envelope (E) gene, with a known number of copies (IDT, USA). Therefore, a concentration curve was prepared by performing serial dilutions of the plasmid.

### 5. RNA libraries and sequencing

For RNAseq analysis, three independent replicates were prepared for each treatment groups: Mock, MOI 0.01- and MOI 0.1-infected cultures, at 6 and 24 hpi. Total RNA was isolated using the miRNeasy micro kit (Qiagen) according to the manufacturer’s instructions. The RNA was quantified by O.D. measurement before being assessed for quality by chip-based capillary electrophoresis using Agilent 2100 Bioanalyzer RNA 6000 Pico assays (Agilent Technologies; Part # 5067-1513).

Libraries were prepared from 150 nanograms (ng) of DNA-free total RNA using the Universal Plus mRNA-Seq Library Prep Kit (NuGEN Technologies, Inc.; Part # 0508-96). The quality and size distribution of the amplified libraries were determined by chip-based capillary electrophoresis on Agilent 2100 Bioanalyzer High Sensitivity DNA assays (Agilent Technologies; Part # 5067-4626). Libraries were quantified using the Takara Library Quantification Kit (Part # 638324). The libraries were pooled at equimolar concentrations and diluted prior to loading onto a P3 flow-cell (Illumina; Part # 20027800) with the P3 300 Cycle reagent kit (Illumina; Part # 20038732) on the NextSeq2000 instrument.

### 6. RNA-seq data analysis

Reads R1 and R2 were trimmed 12 nucleotides (nt) to remove low quality sequences. Bases with a quality score of less than Q20 were trimmed off the right end of each R1 and R2. Illumina adapter sequences were trimmed from the 3’-end of both R1 and R2 reads. Read pairs in which mate in the pair was less than 30nt after trimming were discarded. These quality-filtered reads were then used for alignment.

Sequence alignment was performed using HISAT2 (https://doi.org/10.1038%2Fnmeth.3317) version 2.0.5 with the following settings:

hisat2 --end-to-end -N 1 -L 20 -i S,1,0.5 -D 25 -R 5 --pen-noncansplice 12 --mp 6,3 --sp 3,0 --time --reorder --known-splicesite-infile [SPLICESITES] --novel-splicesite-outfile splicesites.novel.txt --novel-splicesite-infile splicesites.novel.txt -q –x [hsa38 HISAT2 INDEX] −1 [FASTQ1] −2 [FASTQ2] -S [SAMOUT]. The read summarization program featureCounts (https://academic.oup.com/bioinformatics/article/30/7/923/232889) version 1.5.1 was used for exon- and gene-level counting. An Ensembl human version 83 GTF file (downloaded from Ensembl Biomart on January 22, 2016) was used for determination of exon boundaries and the exon-gene relationship during counting. The summarization level used for exon and gene counting was the feature and the meta-feature, respectively. featureCounts is available in the Subread package at http://subread.sourceforge.net/ (https://academic.oup.com/nar/article/47/8/e47/5345150?login=true).

To determine differential gene expression and due the low coefficient of biological variation, paired comparisons were performed between the untreated control and MOIs 0.01 and 0.1 treated HBMEC cells at 6 and 24 hrs timepoints, using an additive linear model with untreated group as the blocking factor. Differential gene expression analysis was performed using EdgeR R package (https://academic.oup.com/bioinformatics/article/26/1/139/182458).

The top differentially expressed genes have consistent UC vs MOIpt1 changes for the three replicates at 5% FDR, and absolute log2 fold change of 0.6 were considered cut-offs to generate the DEG list. Computed z-scores of significant genes are represented in the heatmap. Heatmap was plotted using complexheatmap R package. (https://academic.oup.com/bioinformatics/article/32/18/2847/1743594).

### 7. Downstream RNA-seq analysis

We used a list of genes differentially expressed between MOI 0.1 SARS-CoV-2-infected and uninfected HBMEC cells. The pathway enrichment and interaction networks analysis were performed using clusterProfiler and gprofiler2 R packages (https://www.sciencedirect.com/science/article/pii/S2666675821000667, https://f1000research.com/articles/9-709/v2). Overlapping gene sets from reactome pathway terms, were visualized as a chord plot using the GOplot R (https://academic.oup.com/bioinformatics/article/31/17/2912/184136).

### 8. RT-qPCR

Cells were grown in 60 mm^2^ dishes and total RNA was extracted with Trizol reagent (Thermo Fisher), according to the manufacturer’s instructions. One microgram of total RNA was reversely transcribed into cDNA using SuperScript III system (Thermo Fisher) and 0.5 μl of cDNA was used per RT-qPCR reaction with Power SYBR Green (Thermo Fisher) master mix. Reactions were read in 7500 StepOne Plus from the Oswaldo Cruz Institute. Primer sequences for Drp1, Fis1, ZO-1, claudin-5, HIF-1α, Mfn2, MFF and TOMM20 are provided in **Table S1**. For E gene and Spike1 RT-qPCR, we used the protocols described in Corman et al., 2020 and Won et al., 2020, respectively. For the remaining genes, 100 ng of total RNA was used for Taqman reactions using primer probes from Thermo Fisher: Hs00242739_m1 (LTB), Hs00174128_m1 (TNF), Hs00232399_m1 (RELB), Hs00357891_s1 (JUNB), Hs00759776_s1 (FOSL1), Hs00765730_m1 (NFKB1), Hs00174103_m1 (CXCL8), Hs00601975_m1 (CXCL2), Hs00236937_m1 (CXCL1), Hs00173615_m1 (PTX3), Hs00174961_m1 (EDN1), Hs00299953_m1 (SERPINE2) and Hs01028889_g1 (NFKB2). GAPDH (Hs02786624_g1) was used for sample normalization. Gene expression variations were assessed by the 2^ΔΔCt^ method, with Ct as the cycle number at threshold. Desired PCR result specificity was determined based on melting curve evaluation.

### 9. Western blotting

HBMECs were cultivated in 60 mm^2^ dishes and at desired times were washed with PBS and lysed in the presence of 1x Laemmli Buffer (0.0625M Tris, 0.07M SDS, 10% glycerol, 5% β-mercapto-ethanol and bromophenol blue). Protein concentration was measured with BCA Protein Assay Kit according to the manufacturer’s instructions (Thermo Fisher Scientific, Carlsbad, CA, USA). Then, 30 μg of protein were loaded onto 4-20% gradient acrylamide gels (Bio-Rad Laboratories, Hercules, CA, USA). Membranes were blocked with bovine serum album (BSA) 5% in TBS-0.05% Tween20 and incubated overnight at 4°C with the primary antibodies at 1:1,000 dilution in TBST (**Table S2**). Next day, blots were washed with TBS-0.05% Tween20, incubated for 1 hour at room temperature with secondary antibodies (Lincoln, NE, USA) and analyzed using the Licor CLX imaging system and the Image Studio 4.0 software (LI-COR).

### 10. Immunofluorescence

Cells were grown on 13-mm round glass coverslips and fixed at desired times with 4% paraformaldehyde in PBS for 10 min at 20°C, permeabilized with 0.5% Triton x-100 (Sigma Aldrich), blocked with 3% bovine serum albumin (BSA, Sigma Aldrich) and incubated overnight with primary antibodies at 4 °C. Cells were washed with PBS and incubated with fluorescently labeled secondary antibodies for 1 h at 37 °C. For nuclear visualization, cells were incubated with DAPI (4′,6-diamidino-2-phenylindole) and mounted in a solution of glycerol and DABCO (1,4-diazabicyclo[2.2.2]octane) in PBS. The list of primary and secondary antibodies used in this study are detailed in **Table S2**.

### 11. Quantitative analysis of mitochondrial network morphology

Mitochondrial network morphology was analyzed using the Mitochondrial Network Analysis Tool (MiNA) for the Fiji distribution of ImageJ (Valente et al. 2017). Images were cropped into individual cells. To enhance contrast and sharped mitochondrial images, several pre-processing tools were applied to each image prior to MiNA analysis. First, an unsharp mask (sigma=3) was used to sharpen images by subtracting a blurred version of the image (i.e. unsharp mask) from the image. The unsharp mask is created by Gaussian blurring the original image and multiplying the blurred image by the mask weight (0.8). Second, a median filter (radius=1) was applied to each image. The median filter functions by replacing each pixel with the neighborhood median, where the neighborhood size is determined by the radius. Following pre-processing, images underwent thresholding using the Otsu thresholding to produce a binary image (Otsu, 1979). The ***mitochondrial footprint*** is calculated as the total number of mitochondria-signal positive pixels from the binarized image. A morphological skeleton is then produced from the binarized image using the Skeletonize 2D/3D plug-in (Arganda-Carreras et al. 2010; Lee et al. 1994). This method employs iterative thinning to create a skeleton of mitochondrial structures, one pixel wide. Length measurements of the mitochondrial structures are then measured using the Analyze Skeleton plug-in, resulting in two additional parameters: ***mean branch length*** and ***mean summed branch length***. Mitochondrial form branching networks, in which branches intersect at a node. Mean branch length is calculated as the mean length of mitochondrial structure between two nodes. Mean summed branch length is calculated by determining the sum of branch lengths within an independent network structure and dividing by the total number of independent networks within a cell.

### 12. Angiogenesis-related proteins secretome

For generation of HBMEC Conditioned Medium (CM), cells were plated on 6-well plates and after infection were maintained in a total volume of 1 ml per well. Conditioned culture media were collected at 24 hours post infection (hpi) and centrifuged for 5 min at 10,000 rpm at 4 °C and stored at −80 °C until use. Secretion of angiogenesis-related protein levels was detected using a Proteome Profiler™ Human Angiogenesis Antibody Array kit (R&D Systems) according to manufacturer’s instructions. Membranes were incubated with pools of two independent experiments (e.g.: Mock #1 + Mock #2, MOI 0.01 #1 + MOI 0.01 #2, etc.) for each experimental condition, in a total of 3 membranes for 6 biological replicas. Spots were detected with chemoluminescence and X-ray films were exposed for 1, 5, 10 and 15 minutes to detect differentially expressed proteins. Densitometric analysis was performed with UN-SCAN-IT gel analysis software and relative intensity values for each spot of the 1-minute exposed film was analyzed with GraphPad Prism software version 9.0.1.

### 13. Transmission electron microscopy

Cells were grown on 35 mm petri dishes and infected or treated as described above. At desired time, cultures were washed in PBS, fixed with 2.5% glutaraldehyde diluted in 0.1 M cacodylate buffer with 3.5% sucrose and CaCl_2_ for 1 h at 20 ºC, followed by washes in cacodylate buffer and post-fixation with 1% osmium tetroxide with potassium ferricyanide for one hour at 4 ºC in the dark. Cells were dehydrated in crescent acetone gradient and embedded in Epon resin at 60°C for 72 h. Ultrathin sections were obtained with Leica ultramicrotome and collected in 300-mesh copper grids, stained with uranyl acetate and lead citrate and visualized at Hitachi Transmission Electron Microscope at Centro Nacional de Bioimagem (CENABIO-UFRJ).

### 14. Statistical analyses

For RT-qPCR and western blotting, a minimum of 5 independent cell culture preparations were used and analyzed with Two-Way ANOVA with Bonferroni post-test in GraphPad Prism Software v9.3.1. Morphometrical analysis of ZO-1 immunostaining were performed with ImageJ software for fluorescence intensity and Tight Junction Organization Rate (TiJOR) using the TiJOR macro for ImageJ, which in an index of localization of tight junction proteins in membrane-membrane contact region of adjacent cells as described by (Terryn et al. 2013).

## RESULTS

### 1. Characterization of HBMEC infection by SARS-CoV-2

In order to characterize the profile of host cell infectivity by SARS-CoV-2, HBMEC and Vero E6 cells were infected in the presence or not of serine endoprotease TPCK trypsin (1 μg/ml), which was shown to increase infectivity in Calu-3 cells, a permissive cell line for the efficient replication of SARS-CoV-2 (Jiang et al., 2021). Cultures were infected at different multiplicities of infection (MOIs) of 0.01, 0.1, 1 and 2 and supernatants were collected at 6-, 24-, 48- and 72-hours post-infection (hpi) and analyzed by RT-qPCR for quantification of viral E gene (**Figure 1A**). We found that HBMECs showed no increase in viral replication or release in the supernatant over time, whereas Vero E6 cells had time-dependent release of SARS-CoV-2 in the supernatant, as expected (**Figure 1A**). TPCK trypsin treatment did not affect the cell infectivity rates; however, for consistency, all subsequent assays were performed in the presence of TPCK. In the same context, HBMECs infected with different MOIs did not show any increase in the expression of SARS-CoV-2 Spike1 and E genes at 6 and 24 hpi (**Figure 1B**). We showed recently that HBMEC cells express, to some extent, several of SARS-CoV-2 receptors at both RNA and protein levels (Torices et al., 2021). Therefore, we evaluated the possible effect of infection on the expression of ACE2 and TMPRSS2 in HBMEC cells and found that infection with the MOI 0.1 induced a 40% decrease in ACE2 mRNA expression (p<0.05), which did not result in ACE2 protein level alterations (**Figure 1C**). However, TMPRSS2 showed a 1.77-fold increase in protein levels in MOI 0.1-infected cultures at 24 hpi (p=0.078). Despite the apparent non-productive infection of HBMEC, SARS-CoV-2 was able to induce apoptosis of HBMECs, as detected by cleaved caspase-3 immunostaining (**Figure 1D**) at 24 hpi. MOIs 0.01 and 0.1 resulted in 2.27 and 4.1% of caspase-3-positive cells, respectively, whereas non-infected dishes showed a physiological rate of 0.7% of stained cells. Apoptotic stimulus was also observed in Vero E6 cells 24 hpi with MOI 0.1 (**Figure 1D**). Positive control with 0.5 and 2.0 μg Staurosporine for 2 hours led to 1.9 and 9% of caspase-3 positive HBMECs, respectively (not shown).

**Figure 1.**
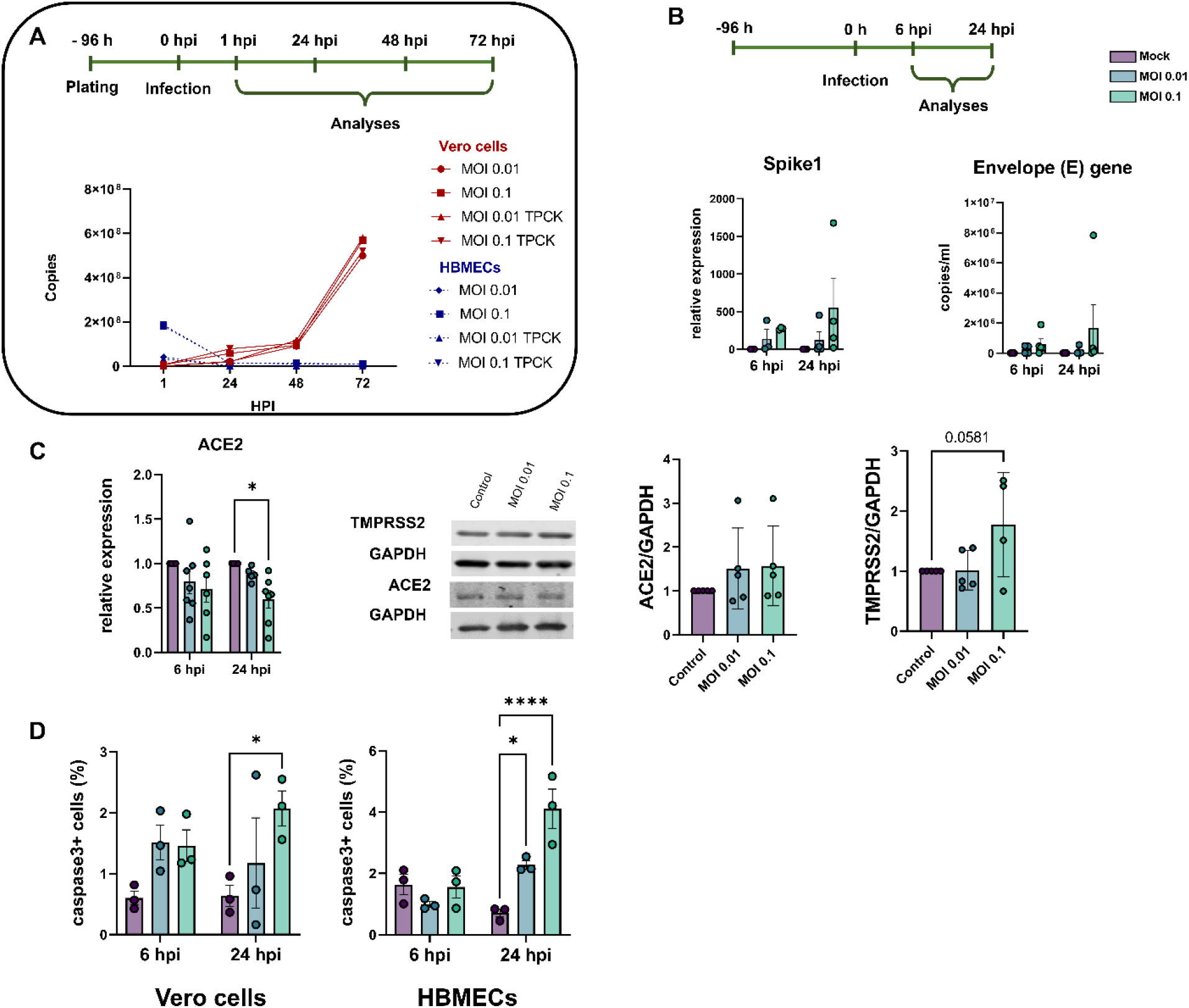
Characterization of infectivity profile of HBMECs by SARS-CoV-2. (**A**) HBMECs were infected with different MOIs of SARS-CoV-2 (variant D614G) and viral production and release to supernatant was analyzed by RT-qPCR for Envelope (E) gene from 0 to 72 hours post infection (hpi). As compared to Vero cells, HBMECs showed a non-productive infection. (**B**) At desired time points (6 and 24 hpi) total RNA from cultures and expression of Spike1 and E genes were analyzed by RT-qPCR. Infection with MOI 0.1 showed an increase in expression of these two transcripts at 24 hpi (p>0.05). (**C**) Evaluation of SARS-CoV-2 receptors expression in infected HBMECs. ACE2 mRNA had a significant decrease at 24 hpi with the MOI 0.1, which did not translate to protein levels (right panel). TMPRSS2 had a slight increase in protein content at 24 hpi. (**D**) SARS-CoV-2 induces apoptotic cell death in Vero and HBME cells at 24 hpi as measured by cleaved caspase-3 immunostaining. *: p<0.05; ****: p<0.0001, Two-Way ANOVA with Bonferroni post-test of at least five independent experiments.

### 2. SARS-CoV-2 affects tight junction genes expression in BBB-forming cells

The barrier property of BMECs is mostly conferred by the expression and function of tight junction proteins, such as ZO-1 and claudin-5 (Takata et al., 2021). HBMEC and Vero E6 cells were infected as described above and analyzed at 6 and 24 hpi. ZO-1 immunoreactivity was drastically altered in infected Vero cultures (**Figure 2A**) and showed discontinuous staining in cell-cell contacts, as compared to uninfected controls. SARS-CoV-2 viral particles were clearly detected in Vero cells as revealed by Spike1 immunoreactivity. Conversely, infected HBMEC cultures did not present as significant differences in distribution of ZO-1 along cells membranes at 24 hpi (**Figure 2A**). To better evaluate ZO-1 organization in TJ we performed densitometric (ZO-1 fluorescence intensity) and tight junction organization rate (TiJOR) (Terryn et al. 2013) analyses. We observed that ZO-1 presented a significant 1.29-fold increase in TiJOR index with MOI 0.1 at 6 hpi, concomitantly with a 1.3-fold increase in fluorescence signal, and such effects were lost 24 hpi. In parallel, MOI 0.01 affected ZO-1 fluorescence signal at 24 hpi by 1.19-fold (**Figure 2B**). ZO-1 and claudin-5 mRNA expression remained unaltered throughout the infection (**Figure 2C**), but their protein levels were significantly increased by 2.0- and 1.17-fold by the MOI 0.1 at 24 hpi, respectively (**Figure 2D**).

**Figure 2.**
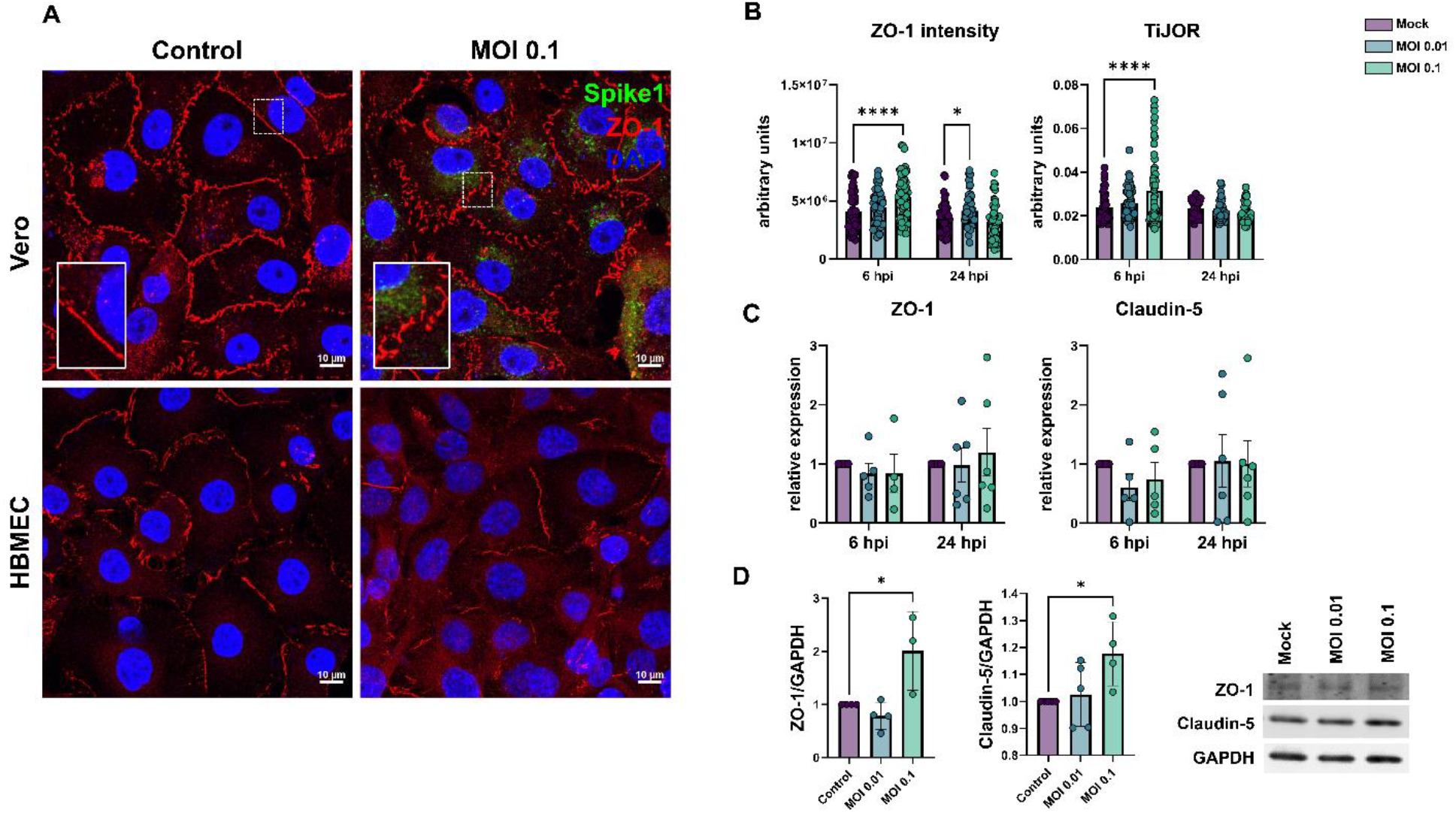
Effects of SARS-CoV-2 on tight junctional proteins in Vero and HBMECs. **A:** Cells were stained for tight junction adaptor protein ZO-1 (red) and SARS-CoV-2 Spike1 (in green). ZO-1 was affected in infected cultures at 24 hpi as shown in higher magnification in the insets. **B**: Morphometrical analyses of ZO-1 fluorescence intensity and TiJOR (D) showed increased ZO-1 signal and TiJOR index at 6 hpi with the MOI 0.01. TJ proteins ZO-1 and Claudin-5 mRNA levels remained unaffected by infection (**C**), but were increased at 24 hpi by MOI 0.1 (**D**). *: p<0.05 One-way ANOVA with Bonferroni post-test (in **D**) or Two-Way ANOVA with Bonferroni post-test (in **B and C**); ****: p<0.0001, Two-Way ANOVA with Bonferroni post-test. Each point in C and D correspond to independent cultures and in **B** correspond to microscopic field from four independent cultures. Representative blots in **D** from three independent experiments.

### 3. SARS-CoV-2 infection promotes endothelial activation and hyper-inflammatory response *in vitro*

We determined the transcriptional profile of HBMECs after SARS-CoV-2 infection by RNA-Seq after 6 and 24 hpi. At 6 hpi, biological replicas had high variability across experiments, as determined by the square root of the common dispersion and visualized by principal component analysis (not shown). Infection with both MOIs 0.01 and 0.1 led to minimal effect on HBMEC’s transcriptome, with few significantly differentially expressed genes (DEGs) and no pathways enrichment found (Supplementary material). However, at 24 hpi we observed significant impact on host cell transcriptome: exposure to SARS-CoV-2 MOI 0.1 led to up-regulation of 23 and down-regulation of 4 genes. Volcano plot and heatmap on **Figure 3** (**A-B**, respectively) depict the transcriptomic profile of HBMECs infected with the MOI 0.1 at 24 hpi. Data obtained from RNA-Seq was consistent with an endothelial activation, with high expression levels of cytokines (IL-6, IL-8, TNF) and chemokines (CXCL1, −2, −8 and CCL20) encoding genes (**Figure 3**). Accordingly, functional enrichment analysis revealed that main genes found related to “Cytokine signaling in immune system”, “TNF signaling” and “TNFR1-induced NFkappaB signaling pathway”, among other Reactome terms (**Figure 3C**). In fact, TNF was the most up-regulated gene, with 104-fold increase, followed by TNF-c (Lymphotoxin beta, LTB), with 32.8-fold change (**Figure 3, Table 1**). Interestingly, LTB is a known inducer of noncanonical NFκB inflammatory pathway (Mockenhaupt et al., 2021) and was found to be up-regulated both at 6 and 24 hpi in SARS-CoV-2 infected HBMEC by RT-qPCR (**Figure 3E**). Although our RNA-Seq data revealed an increase in NFκB2 (p100/p52) and NFκBIA (IκBα), we performed RT-qPCRs with additional biological samples for NF-κB1 (p105/p50) and NF-κB2 and observed that, due to biological variability, such genes remained unaltered in infected cultures (**Figure 3D**). However, RELB, the main activator of noncanonical NF-κB signaling pathway (Mockenhaupt et al., 2021), was shown to be up-regulated by confirmatory RT-qPCR at 24 hpi (**Figure 4**). We also further confirmed by RT-qPCR the up-regulation of inflammation-related genes, including LTB, TNF, IL-6, CXCL1, CXCL2 and CXCL8 (**Figure 3E**). Pentraxin3 (PTX3) is a glycoprotein involved in the innate immune response and has a relevant role in FGF2-dependent angiogenesis (reviewed in Presta et al., 2018). We found that PTX3 to be 19.6-fold increased in SARS-CoV-2 infected HBMECs (**Figures 3A** and **4**). Apart from the inflammatory transcriptomic response, KEGG pathways related to ribosomal structure/function and mitochondrial biology were found to be altered by SARS-CoV-2-infected HBMECs at 24 hpi (**Figure 3D**).

**Table 1.** Differentially expressed genes in MOI 0.1-infected HBMEC cultures at 24 hpi.

| Gene name | Symbol | Fold-change<br>(log) | P value | FDR |
| --- | --- | --- | --- | --- |
| Uncategorized gene | RP11-298I3.3 | -2.19 | 0.00001795234598 | 0.04214061886 |
| ankyrin repeat domain 36C | ANKRD36C | -1.42 | 0.000000005528613831 | 0.00002495701339 |
| upstream transcription factor family member 3 | USF3 | -0.76 | 0.00002036226065 | 0.04595918861 |
| SMG1 nonsense mediated mRNA decay associated PI3K related kinase | SMG1 | -0.75 | 0.000002651386726 | 0.007779698933 |
| nuclear factor kappa B subunit 2 | NFKB2 | 0.64 | 0.00002117775188 | 0.04602945153 |
| E74 like ETS transcription factor 3 | ELF3 | 0.82 | 0.00001027948854 | 0.02601045479 |
| ephrin A1 | EFNA1 | 0.82 | 0.0000001766175517 | 0.0006096837885 |
| Epstein-Barr virus induced 3 | EBI3 | 0.87 | 0.000005771957935 | 0.01612959902 |
| intercellular adhesion molecule 1 | ICAM1 | 0.97 | 0 | 0.0000005593411357 |
| complement C3 | C3 | 1.01 | 0.000000004506547278 | 0.00002203851837 |
| NFKB inhibitor alpha | NFKBIA | 1.05 | 0 | 0.00000008288304584 |
| C-C motif chemokine ligand 2 | CCL2 | 1.14 | 0.000000002315586268 | 0.00001235344223 |
| baculoviral IAP repeat containing 3 | BIRC3 | 1.27 | 0.00000002052506134 | 0.00007528079373 |
| zinc finger CCCH-type containing 12A | ZC3H12A | 1.29 | 0 | 0.0000003145723235 |
| RELB proto-oncogene, NF-kB subunit | RELB | 1.30 | 0 | 0.00000008288304584 |
| Uncharacterized protein | AC010646.3 | 1.31 | 0.0000103901062 | 0.02601045479 |
| interleukin 6 | IL6 | 1.36 | 0 | 0.00000002581261588 |
| TNF alpha induced protein 3 | TNFAIP3 | 1.38 | 0.00000001761165902 | 0.00006890150651 |
| TNF alpha induced protein 2 | TNFAIP2 | 1.42 | 0.000001921435199 | 0.005934605433 |
| C-X-C motif chemokine ligand 2 | CXCL2 | 1.79 | 0 | 0 |
| TNF superfamily member 18 | TNFSF18 | 1.95 | 0.0000106374977 | 0.02601045479 |
| C-X-C motif chemokine ligand 1 | CXCL1 | 2.24 | 0 | 0 |
| C-X-C motif chemokine ligand 3 | CXCL3 | 2.32 | 0.0000000001312489102 | 0.0000007702211048 |
| C-C motif chemokine ligand 20 | CCL20 | 3.50 | 0.000000007991169552 | 0.00003349669957 |
| pentraxin 3 | PTX3 | 4.30 | 0 | 0 |
| lymphotoxin beta | LTB | 5.04 | 0 | 0 |

**Figure 3.**
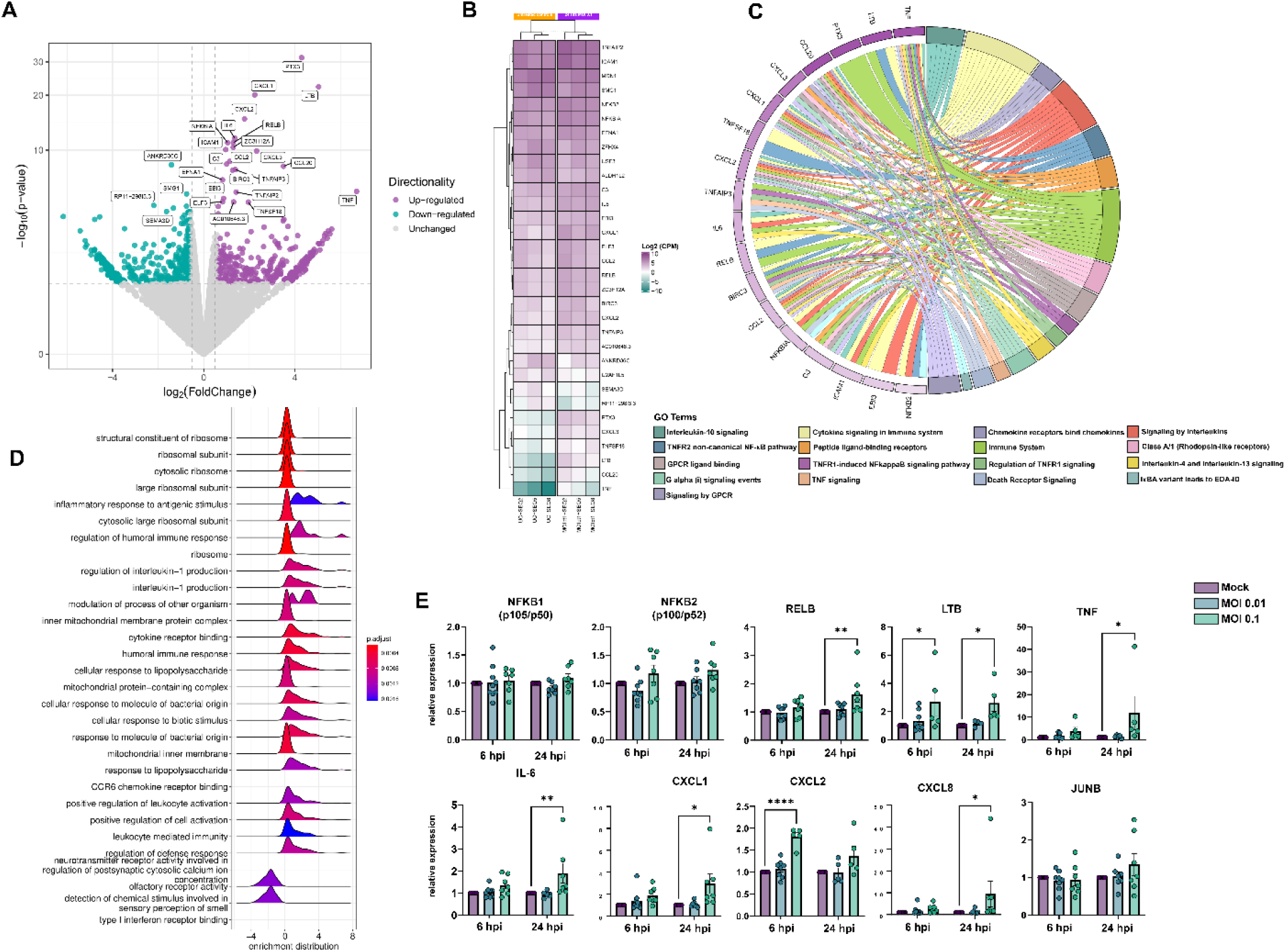
Transcriptomic profiling of SARS-CoV-2 infection on BBB-forming cells. HBMECs were infected with MOIs 0.01 and 0.1 and analyzed by RNA-Seq. (**A**) Volcano plot depicting the overall profile of differentially expressed genes in cultures infected with MOI 0.1 at 24 hpi, with up-regulated genes shown in purple and down-regulated in green. (**B**) Heatmap diagram depicting expression levels of the most significantly altered genes by MOI 0.1 (3 right columns) as compared to uninfected controls (3 left columns). (**C**) Cnetplot visualization of functional enrichment results with up-regulated genes, depicting the functional correlation of genes with the most significant GO terms. (**D**) Enrichment functional analysis of GO terms most affected by SARS-CoV-2 in HBMECs indicate inflammatory endothelial activation, as well as mitochondrial dysfunction and ribosomal-related gene expression. (**E**) RT-qPCR validation of most significantly altered genes detected in the RNA-Seq indicate activation of non-canonical NF-κB pathway, with massive increase in TNF-α, lymphotoxin B (LTB, or TNF-C) and downstream target genes such as IL-6, CXCL1, −2 and −8. NFKB1 (p105/p50) and NFKB2 (p100/p52) as well as JUNB did not show alteration in infected cultures. *: p<0.05; **: p<0.01; ****: p<0.0001, Two-Way ANOVA with Bonferroni post-test of at least 5 independent experiments. MOI: multiplicity of infection; GO: gene ontology.

**Figure 4.**
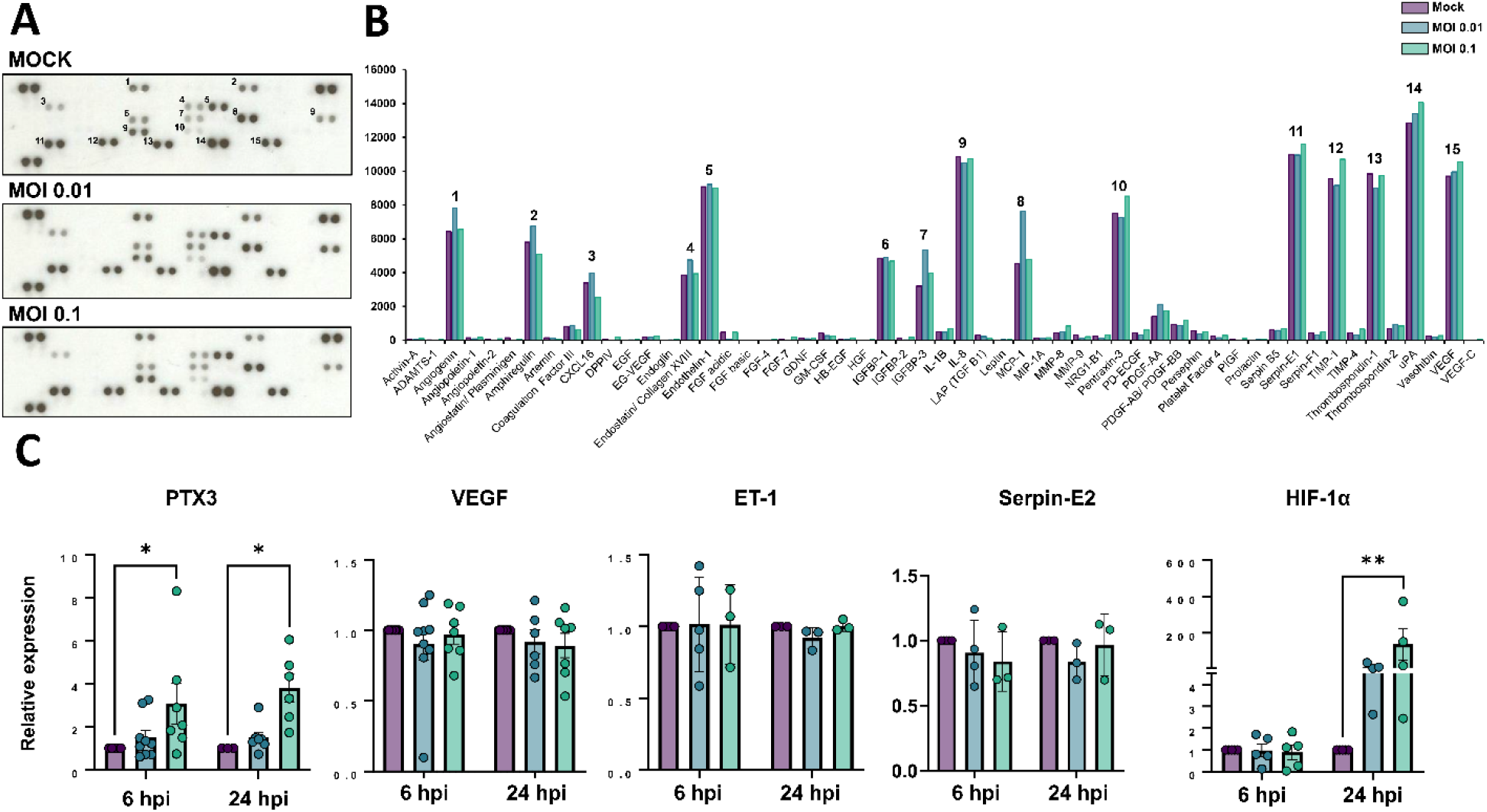
Production of angiogenic-related molecules is modulated by SARS-CoV-2 in HBMECs. (**A**) Conditioned medium from Mock and SARS-CoV-2-infected HBMEC cultures (both with MOI 0.01 and 0.1) were analyzed with Proteome Profiler Human Angiogenic Antibody Array and detected by chemoluminescence, each protein detected in duplicated spots. (**B**) Densitometric analysis of membranes in **A** revealed the analytes with strongest signal and which were affected by infection. (**C**) RT-qPCR analysis of angiogenesis-related genes in HBMECs revealed that PTX3 and HIF-1α were increased in infected cultures. *: p<0.05; **: p<0.01, Two-Way ANOVA with Bonferroni post-test of at least five independent experiments.

### 4. Angiogenic profiling of SARS-CoV-2-infected HBMEC cells

Dysfunctional angiogenesis is a common phenomenon observed in neuroinflammatory states and can be a result of BBB damage (Estato et al., 2018). We analyzed the profile of angiogenesis-related secreted proteins by HBMECs during infection with SARS-CoV-2 and observed that out of 55 spotted targets, 15 had most significantly detectable signals (**Figure 4A-B, Table 2**). Highest signals were observed for uPA, serpin-E1 (PAI-1), IL-8, thrombospondin-1, VEGF, TIMP-1, endothelin-1 (ET-1), PTX3, angiogenin and amphiregulin, with at least 5,000 pixels each. SARS-CoV-2-infected cultures (MOI 0.1) had most pronounced increase in secretion of PTX-3 and TIMP-1 as compared to Mock cultures, with 113 and 112% levels, respectively. Accordingly, PTX-3 was also one of the most up-regulated genes as determined by RNA-Seq (**Figure 3**). We further assessed the expression levels of PTX3 by RT-qPCR and found it to be increased in HBMEC cultures after 6 and 24 hpi with the MOI 0.1, whereas VEGF, Serpin-E2 and ET-1 showed no significant changes at the transcriptional level (**Figure 4C**). Insulin-like growth factor (IGF) binding protein-3 (IGFBP-3), a member of the IGFBP family was shown to be 166 and 125% more abundant in the HBMEC conditioned media in MOI 0.01 and 0.1-infected dishes, respectively. We performed scratch-wound healing migration assays in infected HBMECs, however no effect in cellular migration was noticed in infected cultures as compared to Mock-infected (not shown). Interestingly, hypoxia-inducible factor-1 alpha (HIF-1α) was also increased by SARS-CoV-2 at both MOIs at 24 hpi (**Figure 4C**).

**Table 2.**
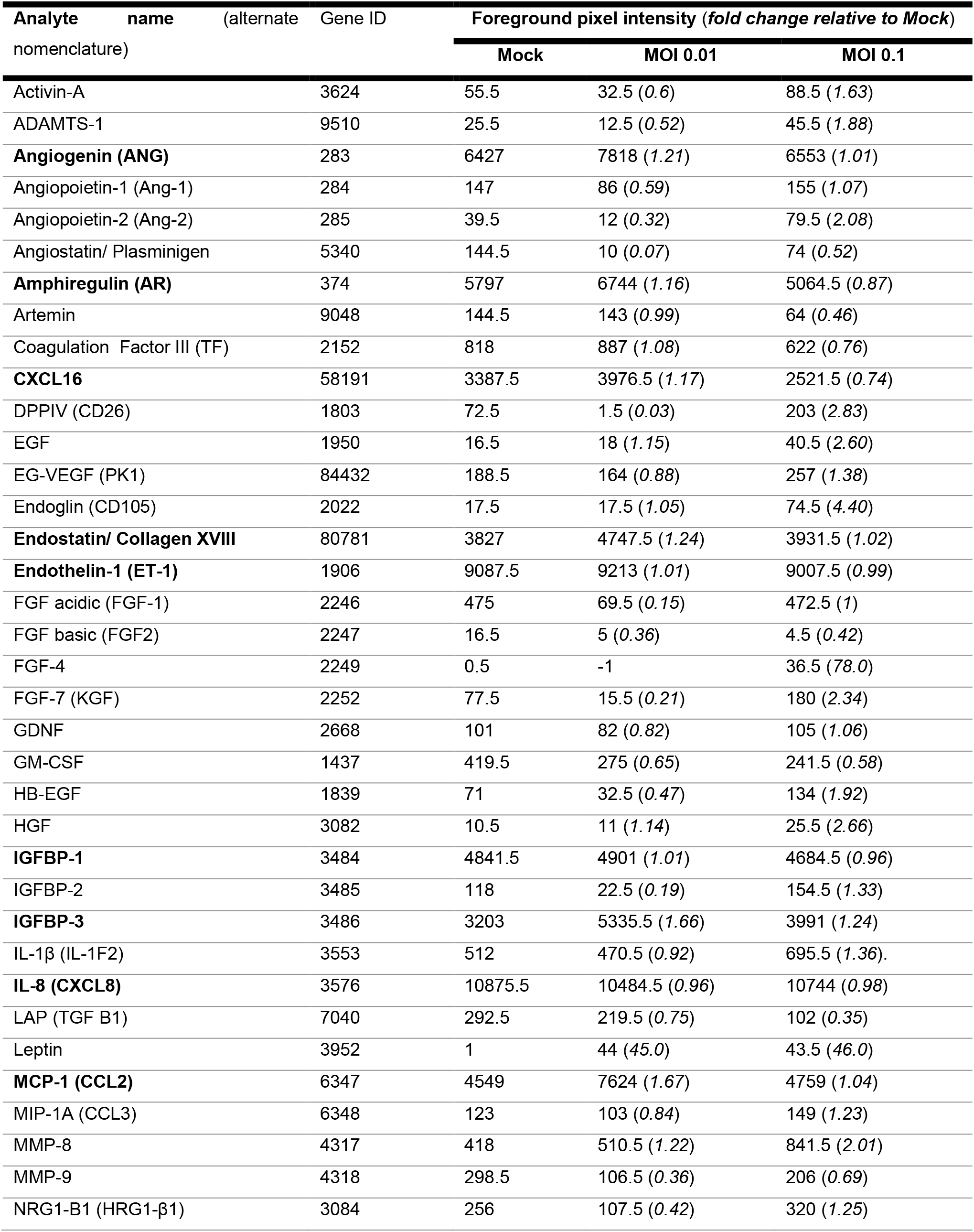
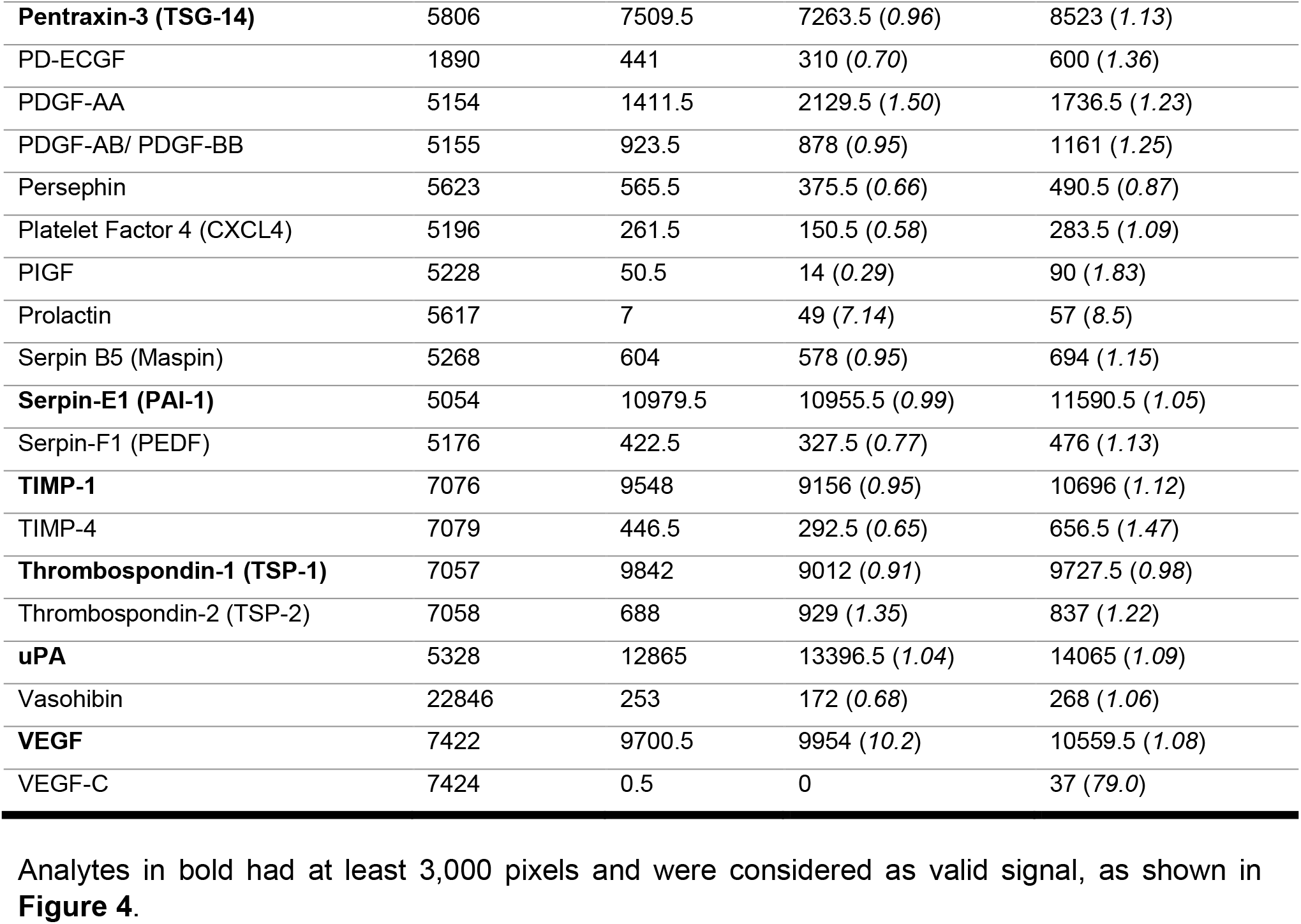
Differentially secreted angiogenic proteins in SARS-CoV-2-infected HBMECs.

| Analyte name<br>(alternate nomenclature) | Gene ID | Foreground pixel intensity (*fold change relative to Mock*) |  |  |
| --- | --- | --- | --- | --- |
|  |  | Mock | MOI 0.01 | MOI 0.1 |
| Activin-A | 3624 | 55.5 | 32.5 (0.6) | 88.5 (1.63) |
| ADAMTS-1 | 9510 | 25.5 | 12.5 (0.52) | 45.5 (1.88) |
| **Angiogenin (ANG)** | 283 | 6427 | 7818 (1.21) | 6553 (1.01) |
| Angiopoietin-1 (Ang-1) | 284 | 147 | 86 (0.59) | 155 (1.07) |
| Angiopoietin-2 (Ang-2) | 285 | 39.5 | 12 (0.32) | 79.5 (2.08) |
| Angiostatin/ Plasminogen | 5340 | 144.5 | 10 (0.07) | 74 (0.52) |
| **Amphiregulin (AR)** | 374 | 5797 | 6744 (1.16) | 5064.5 (0.87) |
| Artemin | 9048 | 144.5 | 143 (0.99) | 64 (0.46) |
| Coagulation Factor III (TF) | 2152 | 818 | 887 (1.08) | 622 (0.76) |
| **CXCL16** | 58191 | 3387.5 | 3976.5 (1.17) | 2521.5 (0.74) |
| DPPIV (CD26) | 1803 | 72.5 | 1.5 (0.03) | 203 (2.83) |
| EGF | 1950 | 16.5 | 18 (1.15) | 40.5 (2.60) |
| EG-VEGF (PK1) | 84432 | 188.5 | 164 (0.88) | 257 (1.38) |
| Endoglin (CD105) | 2022 | 17.5 | 17.5 (1.05) | 74.5 (4.40) |
| **Endostatin/ Collagen XVIII** | 80781 | 3827 | 4747.5 (1.24) | 3931.5 (1.02) |
| **Endothelin-1 (ET-1)** | 1906 | 9087.5 | 9213 (1.01) | 9007.5 (0.99) |
| FGF acidic (FGF-1) | 2246 | 475 | 69.5 (0.15) | 472.5 (1) |
| FGF basic (FGF2) | 2247 | 16.5 | 5 (0.36) | 4.5 (0.42) |
| FGF-4 | 2249 | 0.5 | -1 | 36.5 (78.0) |
| FGF-7 (KGF) | 2252 | 77.5 | 15.5 (0.21) | 180 (2.34) |
| GDNF | 2668 | 101 | 82 (0.82) | 105 (1.06) |
| GM-CSF | 1437 | 419.5 | 275 (0.65) | 241.5 (0.58) |
| HB-EGF | 1839 | 71 | 32.5 (0.47) | 134 (1.92) |
| HGF | 3082 | 10.5 | 11 (1.14) | 25.5 (2.66) |
| **IGFBP-1** | 3484 | 4841.5 | 4901 (1.01) | 4684.5 (0.96) |
| IGFBP-2 | 3485 | 118 | 22.5 (0.19) | 154.5 (1.33) |
| **IGFBP-3** | 3486 | 3203 | 5335.5 (1.66) | 3991 (1.24) |
| IL-1β (IL-1F2) | 3553 | 512 | 470.5 (0.92) | 695.5 (1.36) |
| **IL-8 (CXCL8)** | 3576 | 10875.5 | 10484.5 (0.96) | 10744 (0.98) |
| LAP (TGF B1) | 7040 | 292.5 | 219.5 (0.75) | 102 (0.35) |
| Leptin | 3952 | 1 | 44 (45.0) | 43.5 (46.0) |
| **MCP-1 (CCL2)** | 6347 | 4549 | 7624 (1.67) | 4759 (1.04) |
| MIP-1A (CCL3) | 6348 | 123 | 103 (0.84) | 149 (1.23) |
| MMP-8 | 4317 | 418 | 510.5 (1.22) | 841.5 (2.01) |
| MMP-9 | 4318 | 298.5 | 106.5 (0.36) | 206 (0.69) |
| NRG1-B1 (HRG1-β1) | 3084 | 256 | 107.5 (0.42) | 320 (1.25) |
| **Pentraxin-3 (TSG-14)** | 5806 | 7509.5 | 7263.5 (0.96) | 8523 (1.13) |
| PD-ECGF | 1890 | 441 | 310 (0.70) | 600 (1.36) |
| PDGF-AA | 5154 | 1411.5 | 2129.5 (1.50) | 1736.5 (1.23) |
| PDGF-AB/ PDGF-BB | 5155 | 923.5 | 878 (0.95) | 1161 (1.25) |
| Persephin | 5623 | 565.5 | 375.5 (0.66) | 490.5 (0.87) |
| Platelet Factor 4 (CXCL4) | 5196 | 261.5 | 150.5 (0.58) | 283.5 (1.09) |
| PIGF | 5228 | 50.5 | 14 (0.29) | 90 (1.83) |
| Prolactin | 5617 | 7 | 49 (7.14) | 57 (8.5) |
| Serpin B5 (Maspin) | 5268 | 604 | 578 (0.95) | 694 (1.15) |
| **Serpin-E1 (PAI-1)** | 5054 | 10979.5 | 10955.5 (0.99) | 11590.5 (1.05) |
| Serpin-F1 (PEDF) | 5176 | 422.5 | 327.5 (0.77) | 476 (1.13) |
| **TIMP-1** | 7076 | 9548 | 9156 (0.95) | 10696 (1.12) |
| TIMP-4 | 7079 | 446.5 | 292.5 (0.65) | 656.5 (1.47) |
| **Thrombospondin-1 (TSP-1)** | 7057 | 9842 | 9012 (0.91) | 9727.5 (0.98) |
| Thrombospondin-2 (TSP-2) | 7058 | 688 | 929 (1.35) | 837 (1.22) |
| **uPA** | 5328 | 12865 | 13396.5 (1.04) | 14065 (1.09) |
| Vasohibin | 22846 | 253 | 172 (0.68) | 268 (1.06) |
| **VEGF** | 7422 | 9700.5 | 9954 (10.2) | 10559.5 (1.08) |
| VEGF-C | 7424 | 0.5 | 0 | 37 (79.0) |

Analytes in bold had at least 3,000 pixels and were considered as valid signal, as shown in **Figure 4**.

### 5. Mitochondrial plasticity is affected by SARS-CoV-2 infection

Because mitochondria play a role in cellular homeostasis and pathology, we sought to investigate the effects of SARS-CoV-2 infection on mitochondrial plasticity in HBMECs. Cells were immunostained for mitochondrial import receptor subunit TOMM20 (**Figure 5A**). Our first observation was that infection at MOI 0.01 and 0.1 induced a denser mitochondrial network profile, when compared to Mock condition (**Figure 5A-C)**. MiNA analysis of mitochondrial network organization (Valente et al., 2017), revealed that ***mitochondrial footprint*** (**Figure 5C**), which measures the mitochondria signal in a 2-dimensional image of a cell, was found to be significantly increased in HBMECs infected with the MOI 0.1 at 6 hpi and with both MOIs at 24 hpi. We next measured mean mitochondrial ***branch length mean***, which is the average length of mitochondrial structures that are either independent or connected to networks (**Figure 5C**). We observed a slight, yet significant, increase in cells infected with MOI 0.01 at 6 hpi and with MOI 0.1 at 24 hpi, with a 7 and 3% increase, respectively. Furthermore, MiNA analysis revealed that SARS-CoV-2 induced an overall increase in mitochondrial networks, with significant increase in ***summed branch length mean*** values at 6 (34 and 33% increase for MOIs 0.01 and 0.1, respectively) and 24 hpi (38 and 45% increase for MOIs 0.01 and 0.1, respectively). Mitochondrial morphological analyses were further assessed by TEM (**Figure 5B**) and we found that infected HBMECs displayed larger, swollen mitochondria, with reduced cristae and to some extent associated to multivesicular bodies. Moreover, MOI 0.1-infected cultures displayed 356 mitochondria/mm^2^, while Mock-infected cultures had 266 mitochondria/mm^2^ (p<0.05), which corresponded to a 33% increase (**Figure 5D**).

**Figure 5:**
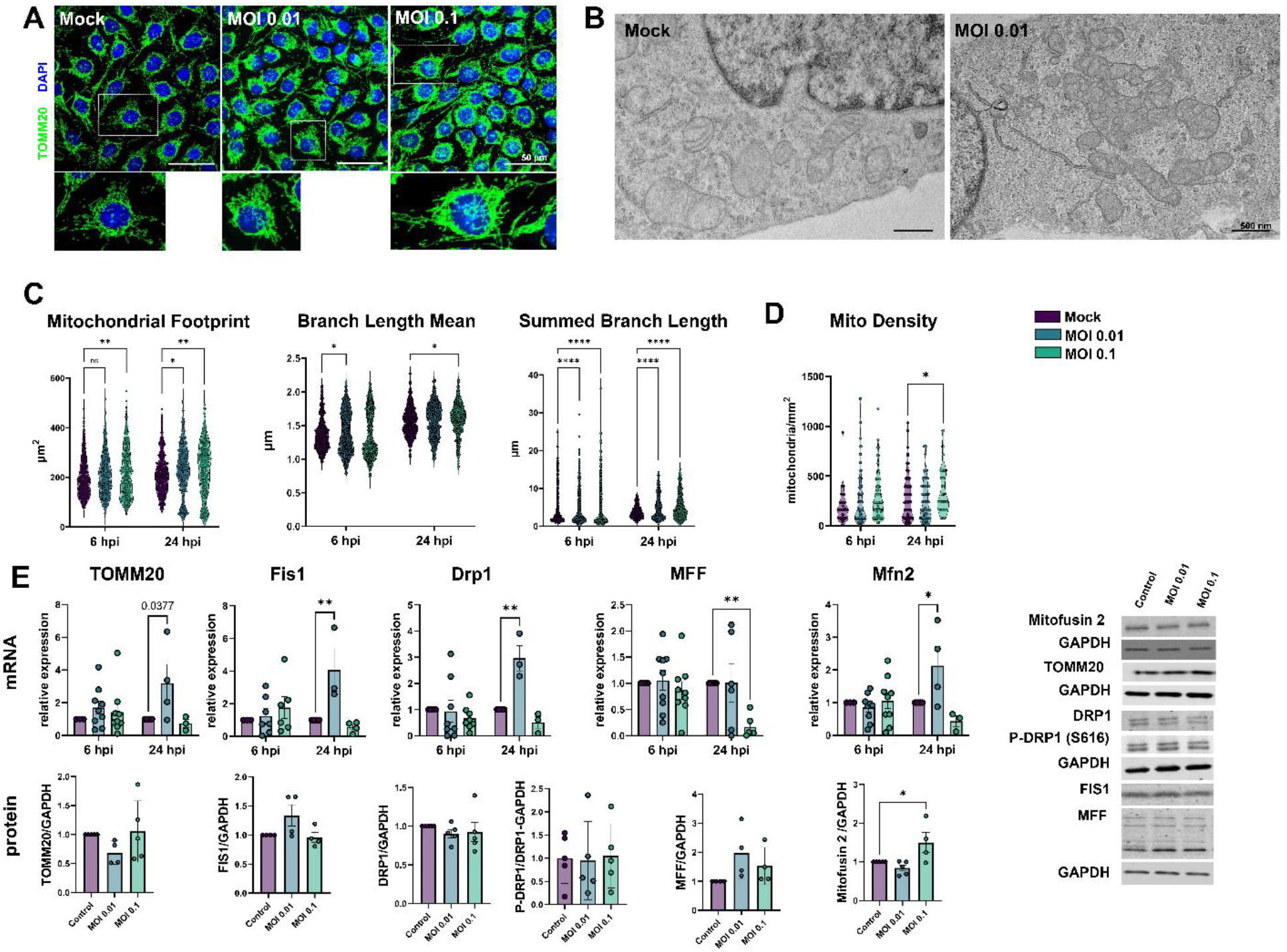
SARS-CoV-2-induced mitochondrial remodeling in HBMECs. Mitochondrial networks were detected by TOMM20 immunostaining (**A**) and TEM (**B**). MiNA analysis of TOMM20 revealed that SARS-CoV-2 induced an increase in mitochondrial footprint, branch length mean and summed branch length (**C**). Mitochondrial density was calculated by TEM images (**D**), which also revealed increased fusion and association to multivesicular bodies (**B**). **E**: RT-qPCR (upper panel) and western blotting (lower panel) analyses revealed that despite fission-related genes (Fis1, Drp1 and MFF) were up-regulated in MOI 0.01-infected cultures, only Mfn2 protein levels were increased in MOI 0.1-infected cultures. TOMM20 protein levels also remained unaltered. *: p<0.05; **: p<0.01; ***:p<0.001; ****: p<0.0001, Two-way ANOVA with Bonferroni post-test. Each dot in graphs represents one cell (in C), one mitochondrion (in D) or one independent experiment (in E). Bottom right panels depict blots from E. Scale bars: 50 μm for **A** and 500 nm for **B**.

Since changes in mitochondrial networks could be influenced by abnormal fission or fusion events (Giacomello et al., 2020), we evaluated expression of markers of such processes. TOMM20, used to determine mitochondrial networks by confocal microscopy and MiNA analysis (**Figure 5A** and **C**), had a 4-fold increase in its mRNA level (p>0.05) in MOI 0.01-infected cells. However, no changes were detected in TOMM20 protein levels by western blotting (**Figure 5E**). We then assessed the expression of mitochondrial fission-related genes. Fis1 and Drp1 mRNA were significantly increased by 4-fold and 3-fold, respectively at 24 hpi in HBMECs infected with the MOI 0.01, which did not translate to changes in Fis1 and Drp1 protein content (**Figure 5E**). Drp1 phosphorylation at serine 616 (Drpi1^S616)^ residue, which is responsible for directing to mitochondrial fission (Han et al., 2020), was not altered in SARS-CoV-2-infected cultures. Mitochondrial Fission Factor (MFF) showed a 0.16-fold reduction induced by MOI 0.1 at 24 hpi, that also did not reflect into altered protein expression. However, Mitofusin-2 (Mfn2) mRNA levels showed a 4-fold increase by MOI 0.01 at 24 hpi (p=0.02), while Mfn2 protein levels were increased by 1.5-fold in MOI 0.1-infected cultures as compared to Mock (**Figure 5E**).

## DISCUSSION

Neurological consequences of COVID-19 still pose a relevant puzzle to medical and scientific community. Since its first cases, CNS invasion has been described (Paterson et al., 2020; Sanchez et al., 2021) but the routes and mechanisms by which SARS-CoV-2 gains entry to brain parenchyma remain allusive (Brann et al., 2020). In the present study we have investigated the molecular indices of Neuro-COVID-19 as they are related to brain endothelial cells forming the BBB. Despite our previous observation that primary HBMECs express several receptors for the virus (Torices et al., 2021), we found little to no indication of productive viral replication in the supernatants of HBMECs which is in accordance with previous reports that described that several endothelial cell types are not permissive for SARS-CoV-2 productive infection (McCracken et al., 2021; Nascimento Conde et al., 2021). Krasemann et al., (2022) observed infection of iPS-derived HBMECs but only at MOIs 10 and 100, which are unlikely to have pathological significance.

Exposure to SARS-CoV-2 led to augmentation of ACE2 and TMPRSS2 expression, which is consistent with our previous observations, in which HBMECs were exposed to the S1 subunit of Spike protein (Torices et al., 2021). Despite the apparent lack of productive infection, SARS-CoV-2 induced apoptotic death of HBMECs in similar levels as compared to Vero epithelial cells and to elsewhere described in the literature (Heuberger et al., 2021). In fact, HBMECs have been shown to undergo apoptotic cell death in response to viral infections, including Dengue (Meuren et al., 2021) and Zika (Leda et al., 2019; Mladinich et al., 2021) viruses, followed or not by changes in BBB permeability. These results suggest that interaction of host cells with viral surface proteins may be sufficient to trigger cellular apoptosis induction even in the absence of productive infection.

Tight junction proteins have a crucial role on maintaining BBB integrity and its selective paracellular permeability (reviewed by Takata et al., 2021). In our study, SARS-CoV-2-infected Vero E6 cells showed a marked disorganization of paracellular tight junctions, as shown by ZO-1 immunostaining. Previous studies described that treatment of HBMECs in 2D or 3D cultures with the S1 subunit Spike1 protein led to mislocalization of ZO-1, concomitantly with cytokine secretion (Buzhdygan et al., 2020; DeOre et al., 2021). Not only ZO-1 (Hao et al., 2020), but also β-catenin, cadherin-5 and occludin junctional proteins have recently been shown to be affected by SARS-CoV-2 proteins in HUVECs (Rauti et al., 2021). ZO-1 possesses a PDZ domain, which is responsible for binding and interaction with other proteins (Saras & Heldin, 1996; Giepmans & Moolenaar, 1998). Interestingly, ACE2 possesses a PDZ-binding domain (Dasgupta & Bandyopadhyay, 2021; Caillet-Saguy & Wolff, 2021), and it has been suggested that epitopes of viral proteins, such as 1–60: M1Lys60 and 241–300: Ala240-Glu300 could directly bind to ZO-1 and VCAM-1 PDZ domains, thus suggesting a possible alternative route of CNS entry. We found ZO-1 and claudin-5 protein levels to be increased after 24 h of infection in HBMECs and this increase is consistent with what our group recently demonstrated with S1 treatment (Torices et al., 2021). Whether ZO-1 increase can be directly related to BBB permeability may vary among experimental models and infectious agents. We have described that the Honduras isolate of Zika virus selectively up-regulates ZO-1 expression *in vitro*, while BBB permeability was increased *in vivo* (Leda et al., 2019). In fact, proper BBB functioning relies on the combined expression and localization of ZO-1, −2 and −3, claudin-5 and occludin (Feng et al., 2018). Direct infection with higher MOIs of SARS-CoV-2 or treatment with plasma from COVID-19 patients failed to induce significant increases in permeability in BMECs *in vitro* (Constant et al., 2021). Conversely, using the K18 mouse model and hamster infection, Zhang et al. (2021) showed that SARS-CoV-2 effectively infects and replicates in HBMECs, but leads to no change in BBB permeability and TJ proteins. Interestingly, a massive inoculation of iPS-derived HBMECs (MOIs 10 and 100), showed active viral replication, whereas MOIs 0.1 and 1 described infectivity near to 0.6% cells, which is similar to what we observed herein (Krasemann et al., 2022).

Following the initial characterization of infectivity profile in HBMECs, we sought to characterize the transcriptomic landscape of SARS-CoV-2 infection in BBB-forming cells. We performed RNA-Seq analyses of SARS-CoV-2-infected cultures at 6 and 24 hpi, with MOIs 0.1 and 0.01. Due to the minimal change in the overall host cell transcriptome with most of the experimental conditions used in this study, we focused our subsequent analyses to the MOI 0.1 infection at 24 hpi. The majority of the significantly up-regulated genes corresponded to known endothelial activation pathways, including CXCL1, −2, −3 and CCL20, PTX3, ICAM1 and TNF. Interestingly, the overexpression of the LTB-TNF-RELB ensemble by SARS-CoV-2, provided evidence that activation of the non-canonical NF-κB pathway activation may be taking place. NF-κB is a family of transcription factors that can be activated by several ligands and activates the expression of proinflammatory cytokines and chemokines (Sun et al., 2012). Interestingly, the main protease of SARS-CoV-2 (M^pro^) cleaves a member of the NF-κB family, NEMO, which in turn leads to HBMEC cell death *in vitro* and *in vivo* (Wenzel et al., 2021). RELB can form heterodimers with p50/p105, p52/p100 and p65 (Shih et al., 2012; reviewed by Mockenhaupt et al., 2021), but can also bind to sirtuin1 to direct epigenetic silencing of inflammatory gene expression (Chen et al., 2009; Liu et al., 2011). In fact, among KEGG pathways enriched in our datasets, we found ribosomal structure and function as possible candidates for epigenetic regulation induced by SARS-CoV-2. These observations suggest that host epigenetic factors may be key for the outcome of COVID-19 and/or long COVID-19. Indeed, the promoters of the genes involved in inflammation, including NF-κB, can be demethylated, thereby resulting in an increased expression of interferons (IFNs), possibly leading to “cytokine storm” (Coit et al., 2016). The expression of IL-6, another important player in the so-called cytokine storm occurring in the most severe COVID-19 patients, was also significantly increased in infected HBMEC and it is known to be modulated by methylation of its promoter. It was also observed that oxidative stress induced by viral infections, including SARS-CoV-2 infection, can inhibit the maintenance of DNA methyltransferase DNMT1, thereby aggravating the DNA methylation defects (Li et al., 2014; Perl, 2013; Sawalha et al., 2020). Our preliminary results (not shown) indicate that SARS-CoV-2 infection of HBMEC results in a decrease in DNA methylation, supporting a recent study of genome-wide DNA methylation analysis in peripheral blood of COVID-19 infected individuals, which identified marked epigenetic signatures, such as hypermethylation of IFN-related genes and hypomethylation of inflammatory genes (Corely et al., 2020). Such observations further suggest the involvement of epigenetic regulatory mechanisms in COVID-19 (Mantovani A & Netea, 2020).

As stated above, PTX3 was one of the main hits found in the transcriptomic analyses. Pentraxins are a superfamily of multifunctional proteins with conserved phylogeny (Daigo et al., 2014), divided into 2 groups based on their primary structure: short and long pentraxin, where c-reactive protein and PTX are examples of short and long pentraxins, respectively. PTX3 is expressed in several neural cell types (Muffley et al., 2012; Shindo et al., 2016; Freezer et al., 2017; Siqueira et al., 2018; Wesley et al., 2022) and in endothelial cells can be upregulated by inflammatory stress such as cytokine stimulation (Breviario et al., 1992). We performed a profiling of angiogenesis-related panel in the supernatants of SARS-CoV-2-infected HBMEC and confirmed that PTX3 was increased by infection. PTX3 is known to be produced in high amounts by blood vessels in vascular inflammatory conditions (Fazzini et al., 2001) and inhibits FGF2-dependent angiogenesis (Rusnati et al., 2004; Presta et al., 2018). Pathological vascularization and angiogenesis have been described as a unique comorbidity associated with SARS-CoV-2 infection in the pulmonary endothelium (Meini et al., 2020), including microvascular distortion and increased intussusceptive angiogenesis (Ackerman et al., 2020; Mentzer et al., 2022). Moreover, VEGF as well as other angiogenic-related analytes were found to be increased in COVID-19 patients (including PTX3), which correlated with disease severity (Maldonado et al., 2022). Accordingly, VEGF levels were 8% increased in the supernatants of infected HBMECs, even though *vegf* transcripts remained unaltered. It is well-known that inflammation, especially IL-6-dependent, can stimulate defective angiogenesis (Gopinathan et al., 2015) and our data further contributes to the notion that following SARS-CoV-2 infection there is an intense brain endothelial activation which leads to defective angiogenic signaling and possibly endothelial permeability. Additionally, we found HIF-1α to be greatly increased after 24 hpi. HIF-1α is a major angiogenesis inductor and is known to be up-regulated by distinct viral infections (reviewed by Reyes et al., 2020). HIF-1α is activated and translocated to the nucleus upon hypoxic conditions (Ke & Costa, 2006) and it has been shown that COVID-19 patients present massive hypoxia due to vasoconstriction and coagulopathy (Afsar et al., 2020). Interestingly, ACE2 expression is decreased in pulmonary smooth muscle cells upon HIF-1α accumulation (Zhang et al., 2020), whereas hypoxia leads to a biphasic modulation of both ACE2 and TMPRSS2 expression on brain microvascular endothelial cells (hCMEC/D3), with an initial increase at 6 h and a decrease at 48 h of hypoxic stimulus cells (Imperio et al., 2021). These observations are in accordance with our present data, that ACE2 is decreased while HIF-1α is increased at 24 hpi. Although VEGF is one of the most described downstream targets of HIF-1α activation, apoptotic cell death and IFN-stimulated gene expression are additional targets of HIF-1α activation (Reyes et al., 2020), which can also be dependent on NF-κB signaling pathway (Walmsley et al., 2005). Our data indicates that HIF-1α up-regulation can be a part of a SARS-CoV-2-induced endothelial activation, along with cytokine/chemokine stimulation and NF-κB non-canonical activation.

Our final series of experiments focused on mitochondrial morphology and dynamics in HBMECs following SARS-CoV-2 infection. It is well known that mitochondria are gatekeepers of BBB endothelium physiology and correspond to higher cytoplasmic volume as compared to non-cerebrovascular endothelial cells (Oldendorf et al., 1977; reviewed by Parodi-Rullan et al., 2021). Moreover, mitochondrial function is important for BBB maintenance and integrity (Doll et al., 2015). We first employed a morphological/morphometrical approach to determine the mitochondrial contents and cellular distribution. Herein we demonstrated that direct exposure to SARS-CoV-2 led to a remodeling of mitochondrial networks. By using the MiNA plugin, we verified that infected HBMECs had increased mitochondrial footprint, as an estimation of overall TOMM20 pixel signal. Recent reports have also shown an effect of SARS-CoV-2 and COVID-19 on mitochondrial biology: monocytes isolated from COVID-19 patients display reduced mitochondrial membrane potential and SARS-CoV-2 viral load was positively correlated with generation of ROS (Romão et al., 2022). Importantly, endothelial cells exposed to SARS-CoV-2 Spike1 protein showed decreased tubular and increased fragmented mitochondrial networks *in vitro*, which was accompanied by a decrease in oxygen consumption rate and increase extracellular acidification rate (Lei et al., 2021). Confirming observation was recently described by Domizio et al. (2022), in which pulmonary endothelial cells in a lung-on-a-chip infection model displayed increased mitochondrial networks. Similarly, we demonstrated that mitochondrial networks were increased, as determined by summed branch length analyses, which indicates that SARS-CoV-2-infected HBMECs had longer mitochondrial ramifications. Changes in endothelial mito-morphology are well described in several models of inflammatory diseases and/or aging (Burns et al., 1979; Jendrach et al., 2005; Forrester et al., 2020) and are correlated with abnormalities in mitochondrial quality control system, which in turn can lead to increased ROS production. Mitochondrial quality control encompasses biogenesis, fission, fusion and mitophagy processes, which are essential for its biology and function. We analyzed markers of fusion and fission processes in SARS-CoV-2-infected cultures and determined that while MOI 0.01 led to an increase in fission-related gene expression (Fis1 and Drp1), this effect was not observed in protein levels or phosphorylation. However, mitofusin2 protein contents were found to be increased in MOI 0.1-infected cultures, which could explain the increased values in branch lengths. Moreover, we found mitochondria associated to some extent to multivesicular bodies, which has also been described as another pathway for mitochondrial quality control (Sugiura et al., 2014; Picca et al., 2020). Interestingly, several reports have linked NF-κB mediated inflammation with mitochondrial responses (Nakahira et al., 2011; Zhou et al., 2016; Zhong et al., 2016; Liu et al., 2018), which could indicate that in fact mitochondrial remodeling observed in infected HBMECs could be due to (or lead to) SARS-CoV-2-induced inflammatory response.

## CONCLUSIONS

Despite no active replication or signs of massive infection, SARS-CoV-2 affects expression of ACE2 and TMPRSS2 in brain endothelial cells and leads to a proinflammatory activation, possibly mediated by NF-κB non-canonical pathway activation. These events would result in mitochondrial and tight junction remodeling and endothelial apoptosis. Taken together our data point to a relevant role of SARS-CoV-2 infection on BBB-forming endothelial cells, which could reflect important aspects of clinical observations of neurological and cerebral vascular manifestations of COVID-19.

## ACKNOWLEDGMENTS

Authors would like to thank the National Center for Bioimaging (CENABIO-UFRJ) for the support in the electron and confocal microscopy imaging.

This work was supported by: Fundação Oswaldo Cruz (Edital INOVA COVID-19 - Geração de Conhecimento 2020, grant number 48401984391807, for D.A.); Fundação Carlos Chagas Filho de Amparo à Pesquisa do Rio de Janeiro (FAPERJ, grant numbers E-26/210.247/2020, E-26/211.118/2021 for D.A.). Natural Sciences and Engineering Research Council (NSERC) of Canada Discovery Grant (number RGPIN–2020-05274 for J.A.S.). D.A. and J.S are supported by a special scholarship for young scientists of the Rio de Janeiro State (Jovem Cientista do Nosso Estado, FAPERJ), grant numbers E-26/201.336/2021 (D.A.) and E-26/201.427/2021 (J.S.). M.T. is supported by the National Institutes of Health (NIH) grants MH128022, MH122235, MH072567, HL126559, DA044579, DA039576, DA040537, DA050528, and DA047157. M.M.S. is supported by Conselho Nacional de Desenvolvimento Científico e Tecnológico (CNPq) through the research grants [402457/2020-0], [313403/2018-0], and [441080/2020-0], the INOVA Fiocruz program, through research grant [48400462543257]; FAPERJ research grants [E-26/210.196/2020] and [CNE E-203.074/2017]. J.S.S.C.M. is supported by a Coordenação de Aperfeiçoamento de Pessoal de Nível Superior (CAPES) doctoral grant.

---

**Table S1:**
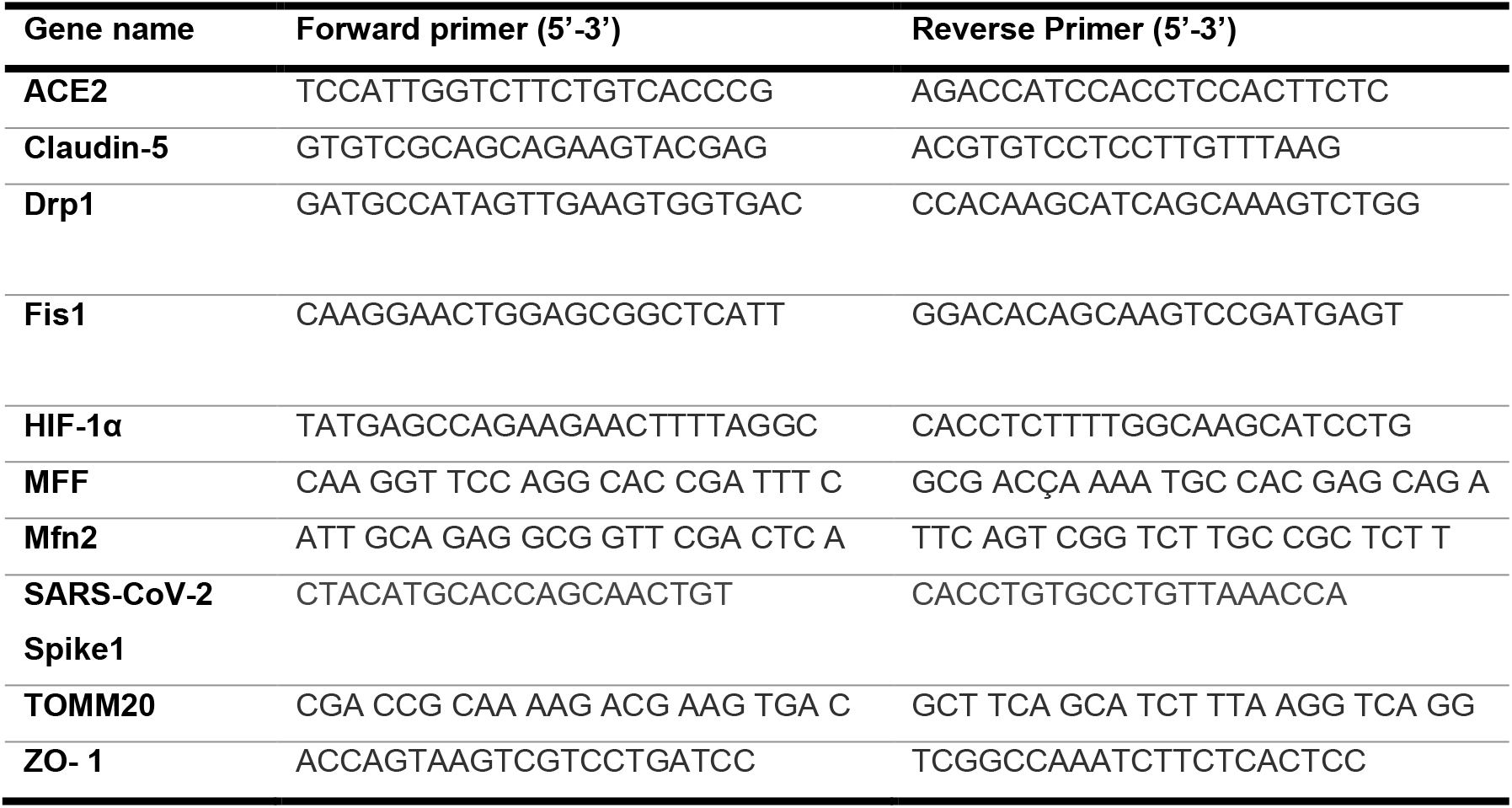
RT-qPCR primer sequences used in this work

**Table S2:**
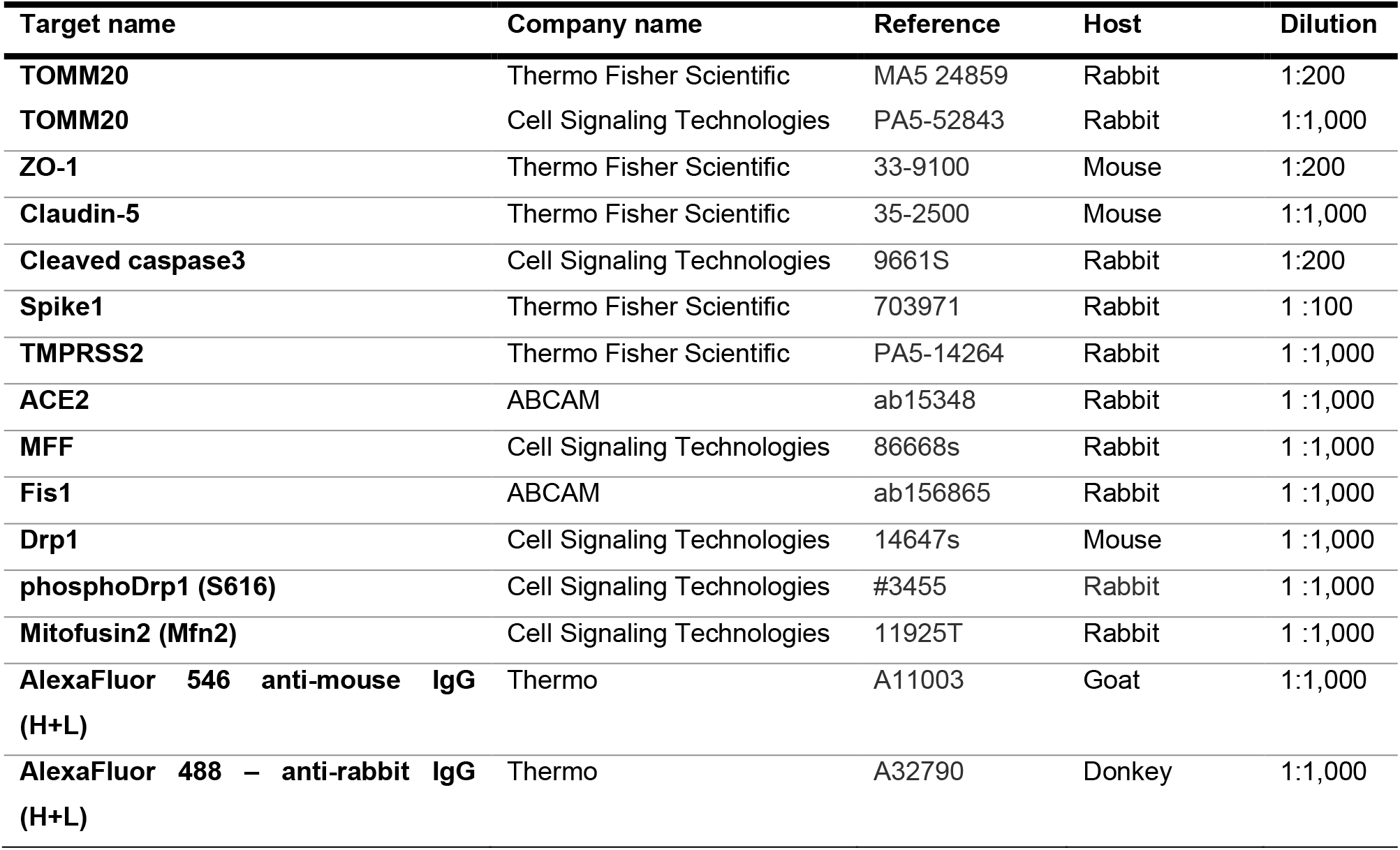
Antibodies used in this work

